# Effect of biogel C addition on biochar degradation and microbial activities

**DOI:** 10.1101/2025.10.13.679449

**Authors:** Prashanth Prasanna, Callum C. Banfield, Bernard Ludwig, Isabel Greenberg, Michaela A. Dippold

## Abstract

Biochar contributes to long-term soil carbon stabilization (C) by acting both as a stable carbon pool and as a sorbent for labile compounds. Biogels from roots and microbes are known to form persistent surface coatings with soil sorbents, where they host microbial hotspots. Yet research has mainly focused on their interactions with minerals. The interactive effects of biogels and biochar on soil carbon dynamics remain largely unexplored. This study aimed to assess the effects of biogel coatings on fresh and aged biochar, particularly under drought, with a focus on biochar degradation and microbial responses. We conducted a three-factorial soil incubation study to examine the effects of fresh and aged biochar, with or without biogel amendment, under two moisture levels (30% and 70% water holding capacity, WHC). Using ^14^*C* labelled biochar, we quantified biochar degradation and biochar-induced priming effects by measuring ^14^*CO*_2_respiration, microbial biomass carbon (MBC) and its ^14^*C* incorporation, hydrolytic and oxidative enzyme activities, and changes in biochar surface area. Mucilage strongly enhanced microbial incorporation of biochar-derived C, specifically from aged biochar under drought conditions, while the opposite effect was observed in soils amended with fresh biochar. This demonstrates the greater microbial accessibility of aged biochar surfaces and limited use of fresh biochar due to inaccessibility of the hydrophobic surfaces, and in consequence the preference for more easily accessible C sources. Additionally, mucilage significantly reduced the Michaelis Menten constant (*K_m_*) of ß-glucosidase by up to 58% in soils amended with aged biochar under drought. This indicates that under these conditions, the interaction of the aged biochar with the biogel may have created a unique habitat requiring specific enzyme systems with higher affinity. Our findings highlight the role of mucilage in regulating microbial surface access and thus the decomposition of biochar, particularly under moisture-limited conditions. The synergy between aged biochar and biogels provides a promising new perspective to further enhance biochar-based drought mitigation, specifically for managing microbial activity in drought-prone soils.

## 1. Introduction

Biochar is a refractory substance which is highly porous, with a network of micro, meso, and macro pores that significantly increases its surface area and enhances its adsorption capacity Leng et al. (2021). The biochar surface contains various functional groups, such as hydroxyl (-OH), carboxyl (-COOH), and carbonyl (C=O) groups, which strongly influence its interaction with other substances Zhang et al. (2021). The surface chemistry of biochar is influenced by multiple factors, including feedstock type, pyrolysis temperature, and production conditions Lehmann and Kleber (2015). Biochar is an effective material for soil remediation and C sequestration, as besides its intrinsic, recalcitrant C input, it stabilizes further C compounds at its surfaces, influences soil nutrient dynamics, and affects microbial activity Gomez et al. (2014). This, in turn, can lead to priming effects Wang et al. (2016), whereby native soil organic C (SOC) or even biochar-derived C is destabilized and decomposed at an accelerated rate. However, it remains unclear whether secondary C stabilization or the priming-induced C destabilization dominates the follow-up processes of biochar-organic interactions. Understanding the mechanisms behind the interactions of biochar with different soil components is essential for predicting its long-term effects on soil C stability. While more is known about the interaction of biochar with low molecular weight organic Dippold et al. (2014);Cheng et al. (2017);Kohlmann et al. (2025) its interaction with more complex organic substances are less studied. Besides polymeric plant tissues, biogels - derived either from microbes or from roots – have gained increasing attention in soil organic matter (SOM) research, as they are an underestimated component of functionally relevant and persistent SOM Redmile-Gordon et al. (2020). These gel-like extracellular polymeric substances (EPS) are composed of polysaccharides jointly with proteins, lipids, and nucleic acids Kidinda et al. (2023). The polysaccharide backbone provides the necessary framework for gel formation and stability Costa et al. (2018). Biogels are known to interact strongly with soil sorbents such as clay minerals and metal oxides Fang et al. (2010), which stabilize the polysaccharide structure and form protective coatings on mineral surfaces. This interaction contributes to the persistence of biogels in soil and are important for soil aggregation, as they enhance cohesion among soil particles Costa et al. (2018);Chen et al. (2023). However, research determining whether the interaction between biogels and biochar surfaces will lead to similar effects is scarce. Given biochar’s high porosity and diverse functional groups, we hypothesize that biogels could adsorb onto biochar surfaces in a similar way as they do with mineral particles Kayoumu et al. (2025). This potential interaction may not only influence the persistence and transformation of biogels in soil environments but could also play a critical role in the long-term stabilization of the biochar itself as well as surrounding SOM. Biochar’s surface properties Janu et al. (2021), explicitly its polarity, strongly determines the capacity to interact with the polar surfaces of biogels, as well as the ability of microbes to colonize its surface. Microbial transformations of the biochar surfaces, the so-called aging, is favored by biochar surface polarity, as microbial movement within water films and adhesion to biochar require a certain level of surface polarity Cheng et al. (2008), but at the same time leads to an increase in polar surface groups. However, a systematic understanding of how aging of biochar affects its interactions with biogels, microorganisms and generally the soil environment is still lacking. Interestingly, biogels themselves can exhibit hydrophobic properties when they are dry Ahmed et al. (2015), which suggests that they may also interact with non-activated, hydrophobic biochar surfaces, specifically under drought, with yet unknown consequences for the fate of biochar in soil. Biogels, such as root mucilage and bacterial EPS, possess an intrinsic ability to absorb and retain water to an amount significantly exceeding their own dry weight Adessi et al. (2018), modulating soil moisture dynamics particularly under drought conditions. Mucilage exuded by plant roots can absorb between 25 to 600 times its dry weight in water McCully and Boyer (1997); Capitani et al. (2013). Similarly, bacterial EPS can retain water up to 15 to 20 times their dry weight Chenu (1993);Adessi et al. (2018), which is vital for maintaining microbial activity under desiccation stress Nazari et al. (2022). Under drought, biochar surfaces covered by hydrated biogels may serve as localized hubs for microbial activity, sustaining essential soil processes Costa et al. (2018). However, we yet lack understanding on the role of the biochar-biogel interaction and the consequences for soil C cycling under drought. Therefore, we designed a three-factorial soil incubation study using aged and fresh biochar, with or without mucilage amendments, under two moisture levels (70% and 30% of water holding capacity (WHC)) to determine the effect of the biochar-biogel interaction, and its implication for water retention and microbial activity, specifically aiming to understand the consequences for the biochar degradation itself. To quantify the biochar decomposition, we used ^14^*C*-enriched biochar, allowing us to quantify the degradation and potential biogel-induced priming of even low amounts of biochar C. Besides testing the effects of biogel- biochar interactions under drought and moist conditions, we aim to specify the effects of the interactions of biogels with fresh (less polar) and aged (more polar) biochar as it is hypothesized to be better coated with biogel carbohydrates, leading to a more rapid degradation of aged than fresh biochar.

## 2. Materials and Methods

### 2.1. Soil Sampling

The soil samples for this study were collected from an arable field in Lüttewitz, Saxony, Germany. The region is characterized by loess coverage, resulting in Haplic Luvisols as the predominant soil type in the field. Soil samples were taken from a depth of 0–10 cm at four randomly selected locations within the field, providing four distinct field replicates. The sampled topsoil is free of *CaCO*_3_ and has the following average characteristics: pH of 5.5, soil organic carbon (SOC) content of 1.2%, total nitrogen (*N_tot_*) 0.12%, clay 17.3%, silt 80.8%, and sand 1.9%. The cultivation on this field has historically included exclusively crops with *C*_3_ carbon fixation, with a recent crop rotation including wheat (Triticum aestivum), sugar beet (Beta vulgaris), and rapeseed (Brassica napus). The soil was sieved to 2 mm and stored at 4 °C for further analysis.

### 2.2. Synthesis of ^14^C biochar

^14^*C*-labelled biochar was produced from corn cobs using maize shoot litter that had been uniformly labelled with ^14^*C*. The maize shoot litter was dried at 60 °C, ball-milled for 1 minute, and then homogenously mixed with shredded corncob granulates in a concrete mixer. The resulting mixture was wrapped in aluminum foil and pyrolysis was carried out in a muffle furnace at 300 °C for 2.5 hours under a continuous nitrogen flow of 5.0 L per minute, ensuring an oxygen-free atmosphere. After pyrolysis, 360 g of biochar were obtained, with a ^14^*C* specific activity of 232 Bq g*^−^*^1^. To ensure particle size uniformity and facilitate homogeneous mixing with soil, the biochar was pressed through a 2-mm sieve Zimmerman et al. (2011). 150 mg of biochar was added to each incubation jar containing 10 g of soil with a total ^14^*C* activity of 30 Bq per jar. This biochar application rate corresponds to 8.7% of the SOC and was thoroughly mixed into the soil before incubation.

### 2.3. Aging of biochar

To produce surface-oxidized biochar, we followed the method described by Mia et al. (2017) treating biochar with a 5% *H*_2_*O*_2_ solution at a 1:30 mass-to-volume (m/v) ratio, thereby heating to 80°C in a water bath for 6 hours under gentle rotation (120–170 rpm). After treatment, the aged biochar was several times thoroughly washed with deionized water to remove residual *H*_2_*O*_2_ and soluble salts, then dried overnight in an oven at 105°C.

### 2.4. Experimental Design and Procedures

Soil samples of 10 g each were prepared with following combinations of amendments: with and without mucilage, without any, with fresh or with aged biochar. These treated samples were incubated in 125-mL bail-lid jars at 20 °C for 85 days under two moisture conditions, namely 30% (drought) and 70% (moist) of the soil’s water holding capacity (WHC). During incubation, *CO*_2_ evolved from the soil was trapped in 3 mL of 1.0 M NaOH solution placed in small vials inside the jars. The NaOH solution, which trapped *CO*_2_ and ^14^*CO*_2_, was sampled and replaced periodically throughout the 85- days incubation (on day 1, 2, 4, 10, 20, 30, 40, 55, 70 and 85). Soil moisture levels were monitored gravimetrically, and additional water was added as needed to maintain the target soil moisture. On day 85, soil samples were harvested and store at 4 °C for further analysis.

### 2.5. Analysis of ^14^Cactivity in CO_2_

The ^14^*C* activity of *CO*_2_ trapped in the NaOH solution was measured by scintillation counting (300 SL, Hidex Oy, Turku, Finland) in 0.5 ml aliquots with 4.5 ml of the scintillation cocktail. To ensure high precision, radioactivity background measurements were taken, and measurement time was set to a minimum of 5 minutes per sample. The total *CO*_2_ trapped in the NaOH solution was measured using a 2100 Total Inorganic C analyzer (Shimadzu, Germany) Bore et al. (2019).

### 2.6. ^14^C activity in MBC and DOC

Soil MBC was extracted using the chloroform fumigation-extraction method Vance et al. (1987). 2.8 gram of 4 °C stored soil sample was weighed in to extraction bottle and extracted with 30 mL of 0.05 M *K*_2_*SO*_4_ solution. Another 2.8 g of soil was fumigated with chloroform for 24 hours in a desiccator to lyse microbial cell membranes before removal of the chloroform by vacuum and thereafter performing the same extraction process. Both extracts were shaken for 1.5 hours and centrifuged at 2000 rpm for 10 minutes. The resulting supernatant was then filtered and stored at -20 °C until analysis. Subsequently, the samples were analysed using a C/N analyzer (Analytik Jena, Jena, Germany). MBC contents and biochar C incorporations were calculated using an extraction coefficient *k_e_C* of 0.45 Wu et al. (1990) for both C and ^14^*C* and are expressed as mg C per gram of dry soil (for total C) and as percentage of added biochar C for ^14^*C* Okolo et al. (2022).

### 2.7. Glucosidase enzyme assay

We used the hydrolytic enzyme *β*-glucosidase as a generic proxy to quantify the microbial processing of labile carbon including the added mucilage. A fluorogenically-labelled substrate, 4-methylumbelliferone-*β*-D-glucopyranoside (MUF, Sigma Aldrich, Germany) was used to determine enzyme activities across a wide range of substrate concentrations, i.e., 0–200 µmol g*^−^*^1^ soil. For the assay, 0.5 g of soil (dry weight equivalent) from the 4 °C stored soil of each treatment was extracted with 50 mL water using low-energy sonication (40 J s*^−^*^1^ output energy) for 2 min. Subsequently, 50 µL of the resulting soil suspensions were added to 100 µL of substrate solution, which included 50 µL MES buffer, and incubated for 2 hours in a 96-well microplate (Puregrade, Germany). Calibration solutions were prepared by mixing 50 µL soil suspension with increasing concentrations of MUF to generate a standard curve. Fluorescence was measured using a Victor3 1420-050 Multi Label Counter (PerkinElmer, USA) at an excitation wavelength of 355 nm and an emission wavelength of 460 nm, with a slit width of 25 nm Razavi et al. (2015). Enzyme activities were expressed as MUF release in nmol per g dry soil per hour (nmol g*^−^*^1^ h*^−^*^1^). Each enzyme assay at each substrate concentration was conducted in triplicate. The kinetic parameters *V_max_* and *K_m_* were calculated by fitting the Michaelis-Menten equation using GraphPad Prism software (Version 10.4.1, 2024, USA).

### 2.8. Oxidative enzyme activities

To analyse the microbial processing of biochar degradation, particularly through oxidative enzymes, we conducted a colorimetric peroxidase enzyme activity assay adapted from Allison and Jastrow (2006) and German et al. (2011). Briefly, 1–2 g of fresh soil was homogenized in 100 mL of 50 mM sodium acetate buffer (pH 5.0) to create a slurry. Aliquots of 200 µL of the slurry were dispensed into 96-well microplates. For peroxidase activity, both 50 µL of L-DOPA and 20 µL of 0.3% hydrogen peroxide were added. Control wells received either buffer or substrate without soil to account for background absorbance. The plates were incubated at room temperature for 4 hours. After incubation, the supernatants were transferred to fresh microplates and absorbance was measured at 460 nm using a Victor3 1420-050 Multi Label Counter (PerkinElmer, USA). Enzyme activities were calculated by correcting absorbance values using the controls and expressed per gram of dry soil, based on an extinction coefficient of 7.9 µmol*^−^*^1^ cm*^−^*^1^.

### 2.9. Surface area of soil-mucilage-biochar mixtures

The specific surface area (SSA) of the samples of the final harvest was quantified with the Micromeritics Gemini VII surface area and porosity analyzer (Micromeritics Instrument Corporation, USA) operating with *N*_2_ as the adsorbate. The instrument was equipped with the VacPrep 061 for sample preparation and was operated with the Gemini VII software (Version 5.01). The measurement was carried out as described in Hagemann et al. (2017).

### 2.10. Statistical analysis

Statistical analyses were carried out with R version 4.5.1 Team (2024). For the incubation experiment, we calculated an analyses of variance (ANOVA) model with fixed effect of moisture (two levels: 30% and 70%), mucilage (two levels: with and without mucilage), biochar type (three levels: fresh, aged and without biochar), including all three two-fold and three-fold interactions. We calculated arithmetic means of the response variables ^14^*CO*_2_ efflux, MBC, *β*-glucosidase, peroxidase activity, and BET and replicates were included via the block effect in the ANOVA. For the ANOVA models, simplifications were carried out, in which first non-significant interaction between the main effect’s moisture, mucilage and biochar were eliminated Crawley (2013). Thus, fixed effects were only considered in the final models in the case of significant contributions. Residuals of the final model for each variable were checked for homoscedasticity graphically and for normal distribution by the Shapiro–Wilk test and graphically by inspecting QQ-plots. The data for ^14^*CO*_2_ efflux, *β*-glucosidase - *V_max_*, peroxidase and biochar surface area was Box-Cox-transformed as no normal distribution was given. Data are presented as arithmetic means and individual replicate values. For significant factors and interactions, the significance level P and the standard error of the difference of the means (SED) is provided.

## 3. Results

### 3.1. Degradation of biochar

The cumulative efflux of ^14^*C*-labelled biochar increased steadily across all treatments throughout the 85 days incubation period (Fig. 1). In the early phase (until Day 4), cumulative ^14^*CO*_2_efflux ranged from 2.6±0.6% to 5.4±2.7% of the applied ^14^*C* with biochar type showing a significant effect on cumulative ^14^*CO*_2_ efflux (p < 0.05, see all ANOVA tables and summary of all modelling results in Appendix 1). Under moist conditions (70% WHC), fresh biochar with or without mucilage exhibited higher ^14^*CO*_2_ efflux (4.2 ± 0.8%) compared to aged biochar with or without mucilage (2.8 ± 0.5%). A similar pattern was observed under drought conditions (30% WHC), where ^14^*CO*_2_ efflux from fresh biochar was higher (5.0 ± 1.9%) compared to aged biochar (3.1 ± 0.4%). Mucilage addition and soil moisture had no significant effect on biochar decomposition in this phase.

**Fig. 1:**
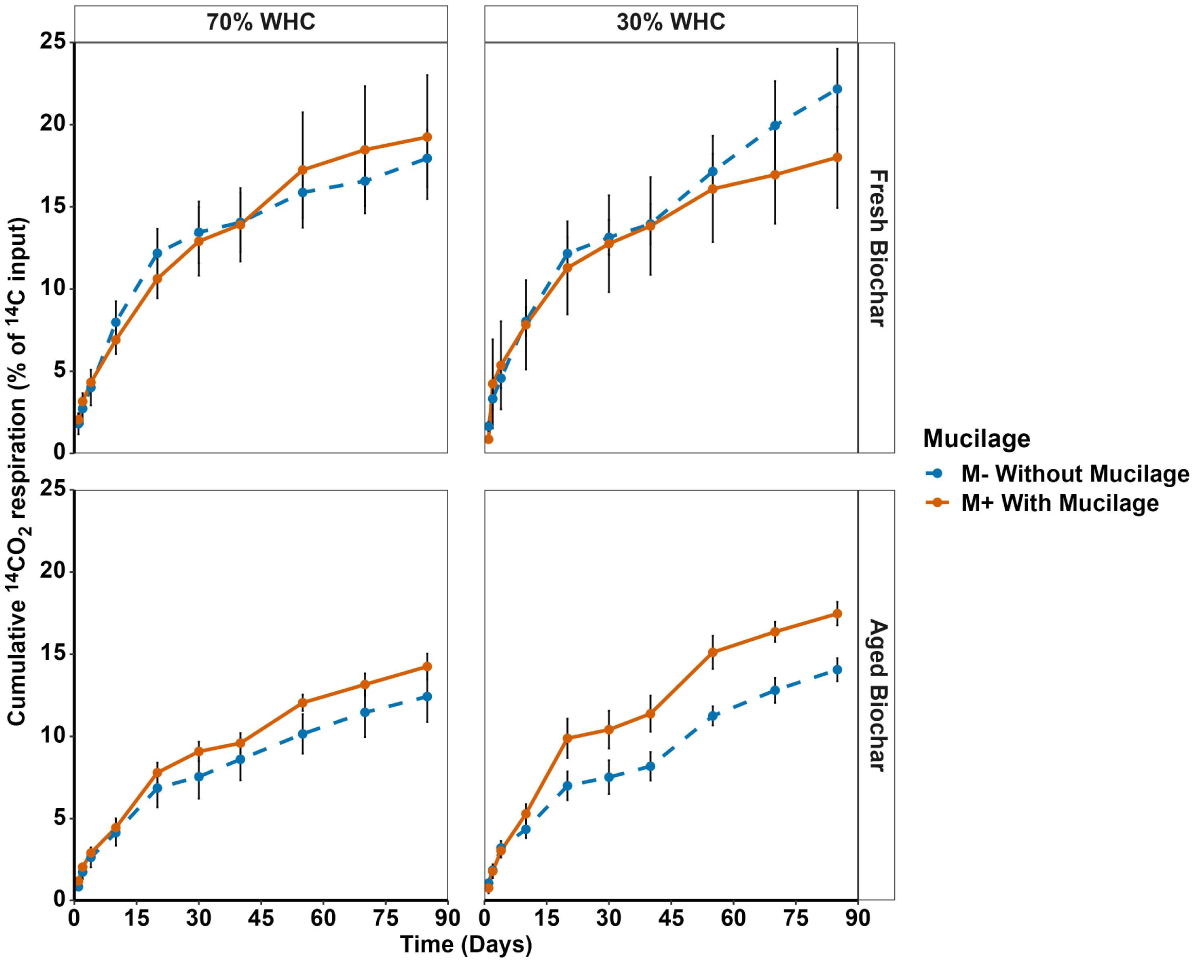
Cumulative ^14^*CO*_2_ efflux (% of ^14^*C* input) over 85 days resulting from the soil incubation under 70% and 30% of soil water holding capacity (WHC), with Fresh and Aged Biochar, and with (M+) and without (M-) mucilage addition. Data represent arithmetic means ± std. errors of the means (n = 4).

In the late phase (Day 85), cumulative ^14^*CO*_2_ efflux ranged from 12.4 ± 1.6% to 22.2 ± 2.5%, with biochar type remaining the only significant factor (p < 0.01). Under moist conditions, ^14^*CO*_2_ efflux from fresh biochar was higher (18.6 ± 2.8%) compared to the aged biochar treatments (13.4 ± 1.2%). Similarly, under drought conditions, fresh biochar again exhibited the highest cumulative ^14^*CO*_2_ efflux (20.1 ± 2.8%) compare to the aged biochar (15.8 ± 0.7%). Although mucilage addition did not show a statistically significant main effect, its presence consistently increased cumulative ^14^*CO*_2_ efflux across both biochar types and moisture levels, particularly in aged biochar under drought conditions.

To isolate the effects of early versus late phase mineralization dynamics, late-phase cumulative ^14^*CO*_2_ efflux was calculated by subtracting cumulative ^14^*CO*_2_ efflux values of day 4 from those of day 85. This late period is assumed to characterize the degradation of the biochar polymer and not volatile or monomeric pyrolysis residues often dominating early degradation patterns. The total cumulative ^14^*CO*_2_ efflux across all the treatments ranged from 9.6% to 18.3% of the applied ^14^*C* biochar (Fig. 2). Biochar type had a significant effect on ^14^*CO*_2_ efflux (p < 0.05), while neither moisture level nor mucilage addition showed significant individual effects. However, the interactions between mucilage and biochar type were weakly significant (p < 0.1). Under both moisture regimes, fresh biochar consistently exhibited higher late-phase ^14^*CO*_2_ efflux than aged biochar. Under moist conditions, ^14^*CO*_2_ efflux from fresh biochar without mucilage was 42% higher compared to the corresponding aged biochar. Mucilage addition increased ^14^*CO*_2_ efflux by 7% from fresh biochar compared to the treatment without mucilage, and by even 16% from aged biochar in moist soil. Under drought conditions, however, fresh biochar treatment without mucilage (17.6 ± 2.0%; M-FB) recorded the highest late-phase ^14^*CO*_2_ efflux, which was 63% higher than the ^14^*CO*_2_ efflux from the aged biochar treatment without mucilage under drought. So, in contrast to in moist soil, mucilage addition decreased ^14^*CO*_2_ efflux from fresh biochar by 28% and from aged biochar by 33% compared to the respective non-mucilage treatments under drought. However, only soil moisture showed a significant main effect in the ANOVA.

**Fig. 2:**
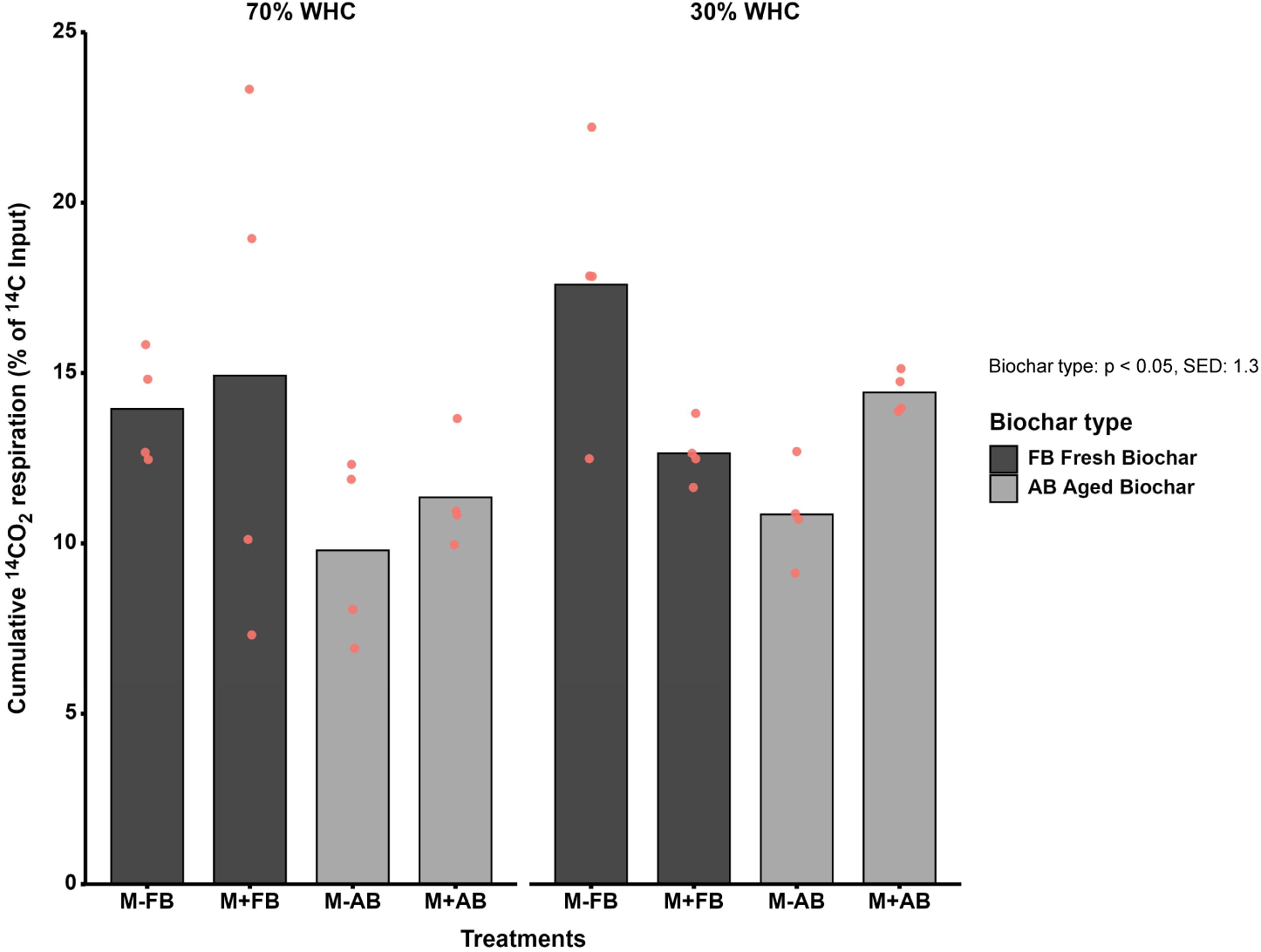
Cumulative ^14^*CO*_2_ efflux (% of total ^14^*C* input) between day 4 and day 85, representing the sustained phase of biochar-C mineralization across treatments with (M+) and without mucilage (M-), Fresh (FB) and Aged Biochar (AB), and 70% and 30% of soil water holding capacity (WHC). Data represent arithmetic means and individual replicates (n = 8). Text box denotes statistically significant effects based on three-way ANOVA and standard error of the differences (SED) of the means for the significant factor biochar type.

### 3.2. MBC and biochar incorporation

Soil MBC was significantly affected by soil moisture (p < 0.01), whereas biochar type and mucilage addition had no significant main or interaction effect on MBC (Fig. 3, top). Across all treatments, MBC was consistently higher in moist soil (70% WHC) compared to drought conditions (30% WHC). Drought reduced MBC by 5% (fresh biochar plus mucilage) to up to 31% (control with only mucilage). This generic reduction in MBC due to drought confirms that moisture is a central driver for microbial abundance irrespective of further factors. The effects of mucilage addition and biochar type on MBC were of minor influence. However, in contrast to the amount of microbial biomass, the incorporation of biochar-derived ^14^*C* into MBC was significantly influenced by mucilage addition (p < 0.05) and by the interaction between moisture, mucilage, and biochar type (p < 0.01). No significant main effect of moisture or biochar type was observed (Fig. 3, bottom). At 70% WHC, incorporation of ^14^*C* into MBC was highest from fresh biochar co-applied with mucilage (1.8 ± 0.3%; M+FB), which was more than twice as high as in the corresponding non-mucilage treatment. However, mucilage addition had a smaller effect on the microbial consumption of aged biochar, resulting in a 20% increase compared to the non-mucilage treatment. Overall, mucilage stimulated microbial assimilation of biochar-derived C more effectively under moist than under drought conditions for fresh biochar, but the opposite moisture effect was observed for the microbial use of aged biochar, which was higher under drought than under moist conditions, representing the soil with the strongest stimulation of biochar consumption by microorganisms. Among mucilage-amended treatments, aged biochar showed significantly higher microbial incorporation than fresh biochar under drought. Overall, mucilage addition enhanced microbial assimilation of biochar-derived C, but its effects were dependent on biochar type and moisture, demonstrating the complexity of the strong interaction across the three experimental factors.

**Fig. 3:**
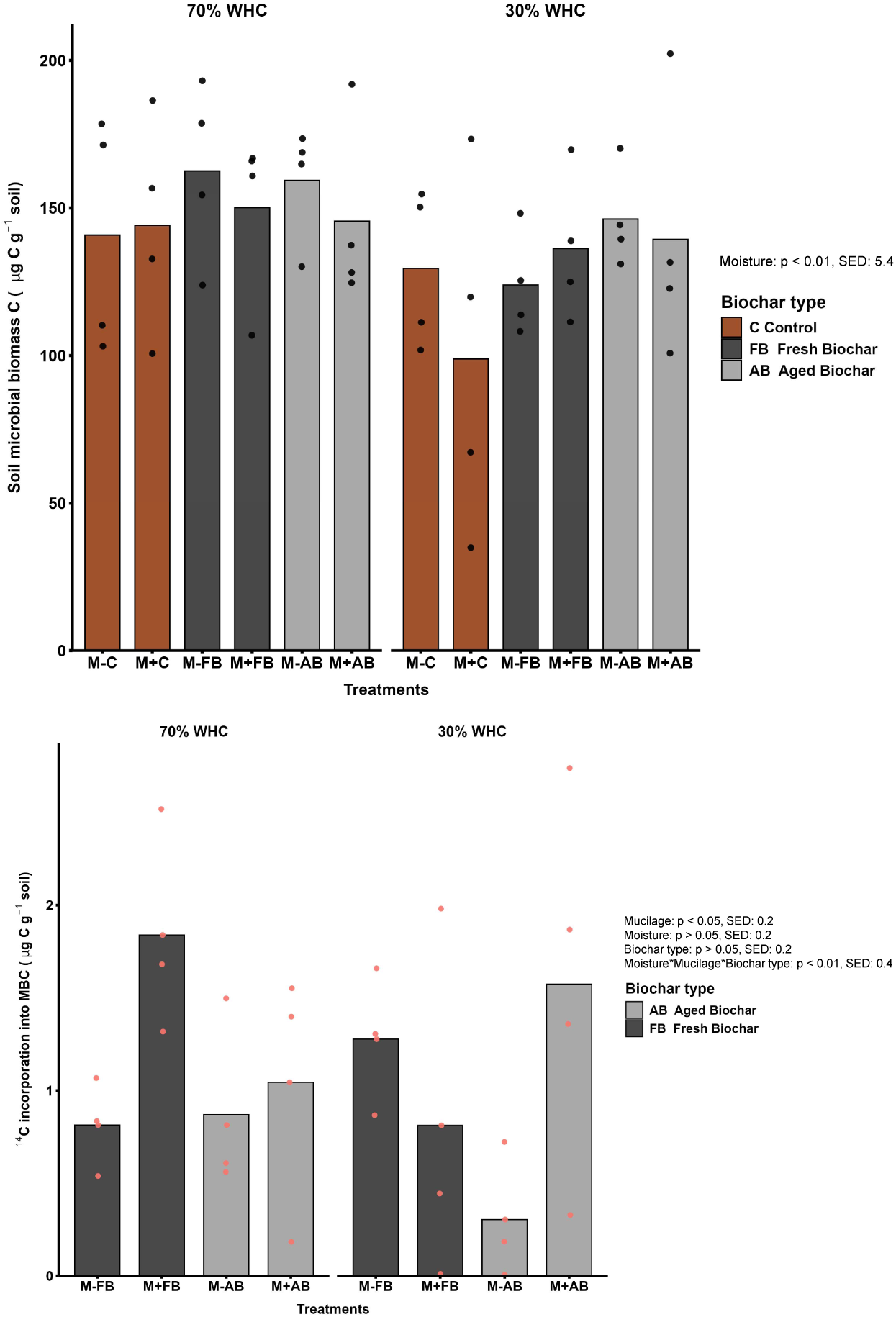
Microbial responses to biochar type, mucilage, and moisture after 85 days of incubation. Top: Soil microbial biomass carbon (MBC; mg C g*^−^*^1^ soil); bottom: Incorporation of ^14^*C* in to MBC (% of total ^14^*C* input). Treatments include with (M+) and without (M-) mucilage addition, Fresh Biochar (FB), Aged Biochar (AB), and no biochar control (C), and 30% and 70% soil water holding capacity (WHC). Data represent arithmetic means (bars) and individual replicates (dots) (n=4). Text box denotes statistically significant main effects and interaction based on three-way ANOVA and standard error of the differences (SED) of the means for the significant factors and interactions.

### 3.3. Microbial activities

The maximum activity (*V_max_*) was significantly affected by soil moisture (p < 0.05). Across all treatments, *V_max_* at 70% WHC was on average 25% higher than that under drought conditions, confirming the strong influence of water availability on the activity of hydrolytic enzymes. In soils with fresh biochar, mucilage addition reduced *V_max_* by 32% compared to the non-mucilage treatment, but for the other biochar treatments, mucilage addition had a negligible effect on *V_max_*. Overall, neither mucilage amendment nor biochar type, nor any interaction among these factors, had a significant effect on *V_max_* of *β*-glucosidase and the activity of this hydrolytic enzymes seems to be mainly controlled by soil moisture. Substrate affinity (*K_m_*) of *β*-glucosidase was significantly influenced by moisture (p < 0.05), mucilage addition (p < 0.05), and their interactions: moisture*mucilage (p < 0.001), moisture*biochar (p < 0.05), and mucilage*biochar (p < 0.05). Under moist conditions, mucilage addition increased *K_m_* across all treatments: in the control treatment by 8%, with fresh biochar by 28%, and with aged biochar even by 37% compared to the treatment without mucilage addition.

This indicates a pronounced reduction in substrate affinity by mucilage addition when biochar was present in moist soil. Under drought conditions (30% WHC), the effect of mucilage addition on the *K_m_* of *β*-glucosidase was contrary to that in moist soil: it led to a general decrease in *K_m_*, i.e., an improved substrate affinity. In the biochar-free control, *K_m_* decreased substantially with addition of mucilage by 47%, while in fresh biochar-amended soil, mucilage addition caused only a slight decrease in *K_m_* for 7%. The strongest response was observed in the aged biochar treatment, where *K_m_* declined by 58% due to mucilage addition. These results indicate that mucilage consistently enhances enzyme-substrate affinity under moisture-limited conditions, an effect strongly promoted under aged biochar amendment, but that this effect is reversed under drought, reflecting again the complexity of the interaction among the three factors and the affinity of the hydrolytic enzyme *β*-glucosidase for its substrate.

Peroxidase activity was significantly affected by biochar type (p < 0.001), the mucilage*biochar interaction (p < 0.05), and the interaction of moisture, mucilage, and biochar (p < 0.05), reflecting similarly to the peroxidase that soil enzymes, hydrolytic as well as oxidative ones, display complex interactions with the soil sorbents (here biochar), the water availability and organic matter components that interacting with water, like the mucilage. Only in moist soils with aged biochar the addition of mucilage resulted in as increase of 39% in peroxidase activity relative to the non-mucilage treatment; otherwise, no stimulation by mucilage addition could be observed. In contrast, in control soils at 30% WHC, mucilage reduced peroxidase activities nearly 3-fold. However, a similar strong mucilage effect was not observed for either of the biochar treatments. Irrespective of soil moisture, peroxidase activities were generally lowest in soils amended with fresh biochar (0.04 µmol min*^−^*^1^mg*^−^*^1^). Overall, biochar presence and type had the strongest effect on peroxidase activity, while mucilage effects were highly moisture-dependent and varied across biochar treatments.

### 3.4. Effect of degradation on inner surface area of soil-biochar-mucilage mixture

The mean surface area of the soil mixtures was significantly affected by incubation time (p < 0.001), showing a consistent increase in inner surface across all treatments from the Day 1 (Harvest 1) to the Day 85 (Harvest 2), while biochar presence and type had no significant effect on the inner surface. While in biochar-free control, soil surface area increased by 15.8%, over the course of incubation of the incubation, it was 25% in the fresh biochar amended soil and 18% in that with aged biochar. Overall, these results indicate that microbial or chemical transformations during incubation resulted in a progressive increase of the inner surface area of the (biochar-amended) soils, independent of the surface properties of the initial biochar applied.

## 4. Disscussion

### 4.1. Effect of mucilage addition on soil microorganisms and biochar degradation

Mucilage addition significantly influenced microbial processing of biochar carbon, evident through enhanced incorporation of biochar-derived ^14^*C* into MBC. Under moist conditions, the microbial incorporation of ^14^*C* into MBC was more than twice as high in fresh biochar co-applied with mucilage than in soils without mucilage addition (Fig. 3, bottom). Although fresh biochar typically resists microbial colonization due to its hydrophobic, aromatic structure Zimmerman et al. (2011), mucilage may enhance accessibility by creating hydrated diffusion pathways and supporting localized microbial activity Nazari et al. (2024); Niedeggen et al. (2024). Under drought, the role of mucilage was more pronounced in soils supplied with aged biochar, where addition of mucilage increased microbial ^14^*C* incorporation into MBC compared to non-mucilage amended soils. This likely reflects the ability of mucilage to retain water and form protective microhabitats on oxidized surfaces, maintaining microbial function despite desiccation Ahmed et al. (2016); Holz et al. (2018). This suggests that mucilage can partially buffer drought stress and enhance microbial activity to even enhance degradation of substances as complex as aged biochar. However, it may not function the same way in the interaction with fresh biochar, where a sufficient water supply and thus hydration of mucilage is a prerequisite to boost biochar C assimilation. While mucilage did not significantly affect *V_max_* of the hydrolytic enzyme *β*-glucosidase, it influenced its *K_m_* significantly, specifically in interaction with biochar and moisture (Fig. 4). Under moist conditions, mucilage addition increased *K_m_*, reflecting reduced enzyme-substrate affinity, likely due to high carbon and thus substrate availability in the moist soil, which reduces microbial carbon limitation and thereby makes high-affinity enzymes unnecessary. Under drought, however, mucilage significantly decreased Km, especially in soil with aged biochar. This suggests that despite the overall dry condition of the soil, microbial activity may take place in hydrated microsites, such as mucilage films on aged biochar surfaces, allowing high-affinity enzymes to overcome the overall drought-induced C limitation Burns et al. (2013); Ahmed et al. (2018). However, future research using advanced imaging methods to demonstrate microbial activity in such hydrated microsites is needed to shed light on this postulated mechanism. Mucilage addition alone had no consistent effect on peroxidase activity, but showed strong interactions with both moisture and biochar (Fig. 5). In moist soils amended with aged biochar, mucilage increased peroxidase activity by 39%, likely due to improved microbial colonization on aged, more polar biochar surfaces, where polar hydrated mucilage could interact well. Conversely, under drought, mucilage addition in control soils led to a sharp decline in peroxidase activity, nearly three-fold lower than without mucilage, indicating that microbes may rather have shifted toward utilizing the easily available mucilage carbon, thereby reducing investment in oxidative enzyme to degrade complex native organic matter. No similar mucilage-driven suppression of peroxidase was observed in biochar treatments under drought, as under these conditions the peroxidase activity was generally low. These findings highlight that mucilage may only modulate microbial oxidative function in the presence of biochar as substrate and sufficient soil moisture allowing accessibility of this substrate. One underlying reason for a remarkable native SOM–mucilage interaction in the control soil under drought may arise from the property of both mucilage and SOM to turn hydrophobic at its surface under drought Ahmed et al. (2016); Benard et al. (2018). This may have resulted in much more pronounced mucilage effects on microbial processes in control soils than in biochar-amended soils, where biochar has a moisture-independent defined polarity depending solely on its aging. However, systematic surface hydrophobicity analyses of the individual soil components depending on soil moisture would be needed to confirm this explanation and identify the hydration threshold when organic matter or mucilage individually and jointly in soil change their surface properties abruptly. Overall, the strongly interactive effects of mucilage with other factors or the contradictory effects of mucilage on different microbial processes led to the absence of any effect of mucilage on biochar-derived *CO*_2_ efflux from moist soil. However, under drought not only microbial incorporation of biochar-derived C into biomass but also its mineralization was affected by mucilage amendment, but depending on biochar-quality: mucilage addition increased mineralization of aged biochar by 33%(Fig. 2), consistent with other positive effects on the microbial activities, which were most-likely mediated by interaction of the polar surfaces of aged biochar and mucilage. In contrast, mucilage suppressed mineralization in fresh biochar, either due to physical pore occlusion or preferential microbial use of mucilage-C Lehmann et al. (2011); Zimmerman et al. (2011).

**Fig. 4:**
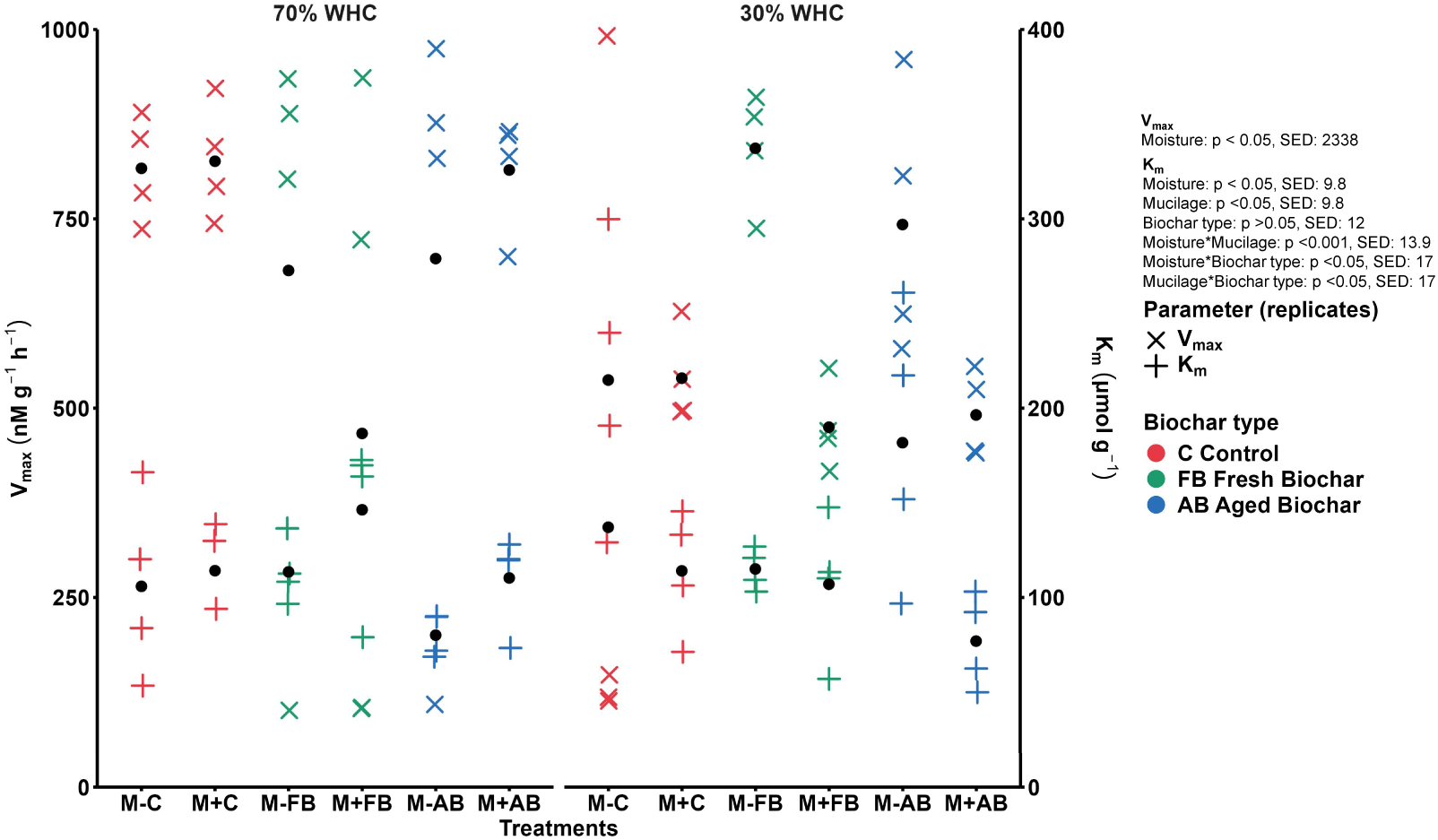
Enzyme kinetics of *β*-glucosidase activity with maximum reaction rate (*V_max_*) and Michaelis constant (*K_m_*) measured under 70% and 30% soil water holding capacity (WHC). Treatments included Fresh Biochar (FB), Aged Biochar (AB), and no biochar control (C), with (M+) and without mucilage addition (M-). *V_max_* is plotted against the left y-axis (X), and *K_m_* against the right y-axis (+). Data represent means (dots) and the underlying replicates (cross and plus for *V_max_* and *K_m_*, respectively) (n = 4). Text box reveals statistically significant effects (including main effects underlying significant interactions) based on a three-way ANOVA and standard error of the differences (SED) of the means for those significant main factors and interactions (as well as their associated non-significant main factors).

**Fig. 5:**
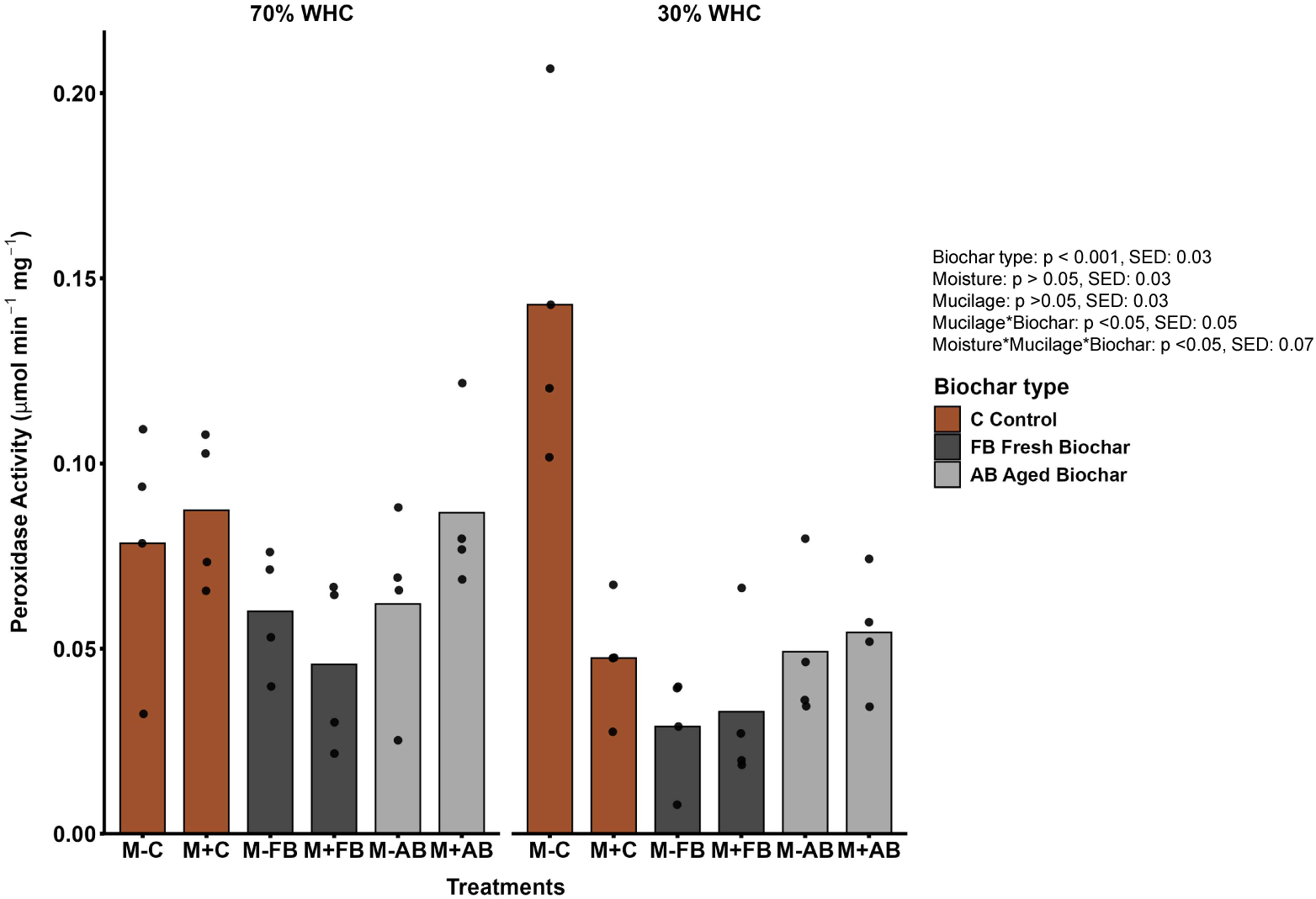
Mean peroxidase activity (µmol h*^−^*^1^ g*^−^*^1^) in soils with Fresh Biochar (FB), Aged Biochar (AB), or no biochar control (C), with (M+) or without mucilage addition (M-), and 70% or 30% soil water holding capacity (WHC). Data represent means (bars) and the individual replicates (dots) (n = 4). Text box denotes statistically significant main factor and interaction effects (and their main effects) based on the three-way ANOVA and SED displays the Standard Error of the Differences of the Means for the significant main factors and interactions (as well as their associated non-significant main factors).

**Fig. 6:**
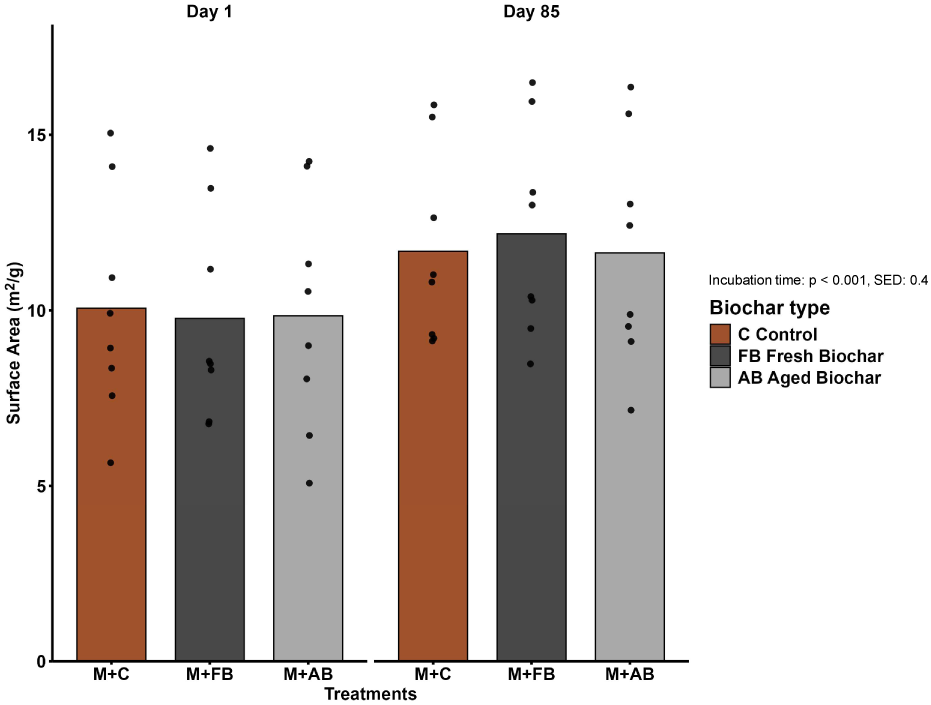
Mean surface area (m^2^/g) of soil mixtures with Fresh Biochar (FB), Aged Biochar (AB), or no biochar control (C), each with mucilage addition (M+), measured at two harvest dates, i.e., day 1 and day 85. Mixtures were incubated at 30% and 70% soil water holding capacity and values were pooled across moisture levels because moisture and its interactions were not significant. Data represent arithmetic means (bars) and individual replicate values (points) (n = 8). Text box denotes statistically significant effects based on two-way ANOVA and standard error of the differences (SED) of the means for the significant main factors and interactions (as well as their associated non-significant main factors).

### 4.2. Effect of soil moisture on microbial activities and biochar degradation

Soil moisture availability emerged as a dominant factor shaping microbial responses and biochar decomposition dynamics. MBC significantly declined under drought (Fig. 3, top), underscoring the fundamental role of water in supporting microbial metabolism, substrate diffusion, and nutrient transport Wang et al. (2016);Joseph et al. (2021). Maximal activity of central, carbohydrate-degrading enzymes (here: *β*-glucosidase) was significantly reduced (ca. 25%) under drought compared to moist soils (Fig. 4), reflecting the critical role of moisture in maintaining extracellular enzymatic activity. Low water content disrupts the spatial connectivity between microbes, enzymes, and substrates, fragmenting the microscale biochemical environment and thus reducing catalytic efficiency Burns et al. (2013);Bogati and Walczak (2022) and influencing substrate affinity, i.e., *K_m_* Munene et al. (2025). Under drought, *K_m_* values were always lower than under moist conditions, which implies that microbial communities under water stress express high-affinity enzymes to optimize substrate utilization when diffusion is constrained German et al. (2011);Tang and Riley (2019). This drought-induced reduction in *K_m_* was strongest (58%) in soils supplied with aged biochar and mucilage, suggesting that the combination of polar biochar surfaces with polar mucilage may best support the activity of enzyme with enhanced substrate affinity in moisture-limited conditions, i.e. prioritizing higher substrate binding over rapid turnover. However, soil moisture neither directly affected the decomposition of biochar, irrespective of age (Fig. 2), nor did it significantly affect the peroxidase activity (Fig. 5). Thus, although soil moisture controls both the magnitude of microbial activity and the kinetic characteristics of enzyme systems degrading carbohydrates, it does not directly affect biochar degradation pathways by oxidative enzymes or its degradation rates. The complexity of the degradative pathway, often occurring as co-metabolism Kuzyakov et al. (2009), and the multifactorial parameters controlling the presence and activity of microorganisms at its surface Lehmann et al. (2011);Palansooriya et al. (2019) may jointly explain the lack of direct dependency on moisture and underlie the complex interactions across our factors.

### 4.3. Effect of biochar type on microorganisms and their activity

Across both the early (Day 4) and late (Day 85) phases, fresh biochar consistently showed higher cumulative ^14^*CO*_2_ efflux than aged biochar, regardless of moisture conditions, underscoring the role of biochar type in regulating its mineralization dynamics (Fig. 1)and(Fig. 2). Unexpectedly, this general effect was not only present in the early phase, in which mostly mono-or- oligomeric (often volatile) pyrolysis products are degraded, but also during late (slower) biochar degradation of more complex structural parts of the biochar. So besides labile (volatile) carbon fractions also surface functional groups of fresh biochar seem to be better amenable to microbial oxidation. In contrast, during aging, these components may have already been degraded, and the biochar is likely enriched in condensed aromatic structures, thus exhibiting reduced ^14^*CO*_2_ release. Mucilage addition interacted with aged biochar, stimulating its mineralization under moist conditions, but suppressing it under drought (Fig. 2), possibly due to mucilage-induced shifts in microhabitat conditions. These results highlight that biochar aging alters decomposition trajectories and the interaction with mucilage. The significant effect of biochar presence and type on peroxidase activity (Fig. 5) (p < 0.001) further confirms the relevance of the char structure on its degradation. Soils with aged biochar showed consistently higher peroxidase activity compared to those with fresh biochar and controls. This suggests that aged biochar is likely characterized by higher redox-active quinone and phenolic groups, which may stimulate the microbial production of oxidative enzymes involved in the breakdown of the complex, non-hydrolysable aromatic compounds German et al. (2011);Zimmerman et al. (2011). This may not be needed for fresh biochar rich in easily degradable surface groups. Together, these observations highlight a clear shift in microbial processing strategies based on biochar aging. Fresh biochar is partially accessible and is used in microbial metabolism getting respired, whereas aged biochar is much more difficult to degrade and thus stimulates enzyme-mediated oxidation, which may not only affect the biochar but also the SOM. These contrasting behaviors reinforce the need to consider biochar aging when describing its degradability and its impact on microbial dynamics, and suggest care should be taken when extrapolating results from short-term experiments with fresh biochar to long-term ecosystem dynamics and impacts of an increasingly aged biochar.

### 4.4. Effects of mucilage interaction with char type and soil moisture on biochar degradation

One remarkable outcome of this study was the dominance of interaction effects among the factors biochar type, moisture and mucilage addition. A number of possible interactions between soil water, biogels and biochar may underlie this observation. Polar, oxidized surfaces of aged biochar facilitate stronger associations with mucilage, enabling formation of hydrated microenvironments favorable for microbial colonization and enzyme activity Cheng et al. (2008);Ahmed et al. (2016), while hydrophobic fresh biochar may repel mucilage, preventing sufficient microbial interaction and possibly diverting microbial activity toward mucilage itself, thereby limiting biochar degradation. Related shifts in microbial habitat properties, may lead to altered microbial enzymatic strategies i.e. high- vs. low-affinity enzymes Burns et al. (2013). Also oxidative enzymes show a nuanced respond to the aromatic C availability and may even profit from mucilage as a co-metabolite, triggering their production to access the stable aromatic carbon of aged biochar Sun et al. (2016). Such interactions reveal that the impact of mucilage addition on biochar-amended soils may be highly context-dependent. While mucilage enhanced microbial colonization, enzyme affinity, and degradation of aged biochar under drought, its addition was neutral or even inhibitory for these microbial activities in soils with fresh biochar. These divergent outcomes reflect that biogels are not solely acting as a labile C sources, but instead function as a soil matrix modifier, influencing microbial access to stabilized organic matter, including biochar Carminati et al. (2017);Ahmed et al. (2018). Whether this feature can be exploited for drought mitigation, specifically in the context of biochar application remains to further be tested at a larger scale.

## 5. Conclusion

While the impact of aged and fresh biochar amendments have been studied in many different formats, and also the physio-chemical effects of biogels on soils are well described, this study aimed for the first time to unravel the interactions of these two key SOM components, as both are intensively discussed in the context of drought mitigation. Specifically with aged biochar, strong interactive effects with mucilage in soils could be demonstrated: it significantly enhanced carbon mineralization and microbial assimilation, but also increased the microbial use of biochar carbon itself. This suggests that mucilage is more than a carbon substrate in these soils and that there is most likely a direct interaction between mucilage and biochar which is not linked to pore clocking i.e. the reduction of accessible surface. One likely explanation is the formation of biofilm-like matrices that sustains microbial activity on hydrophilic biochar surfaces during water scarcity. When exposing such soils to drought, mucilage buffered the decline in microbial activity, maintaining substrate affinity and enzyme activity. In conclusion, the combination of adding or enhancing the production of mucilage combined with the amendment of aged biochar offers a promising strategy to stabilize microbial function under drought. This may be on the cost of enhanced biochar C turnover under moisture limited conditions. Our results have important potential implications for biochar-based soil management practices, particularly in arid or drought-prone environments, where the targeted combination of high-mucilage exuding plants or alternatively the application of hydrogels jointly with biochar with optimized surface properties may maintain microbial functions even during extended drought periods.

## Acknowledgments

We acknowledge German Research Foundation (DFG) for funding the project (LU 583/22-1 and DI 2136/15-1). We thank technicians from the Biogeochemistry of Agroecosystems group at the University of Goettingen and Geo-Biosphere Interactions group at the University of Tuebingen for there technical assistance.

## References

Adessi, A., Cruz de Carvalho, R., De Philippis, R., Branquinho, C., Marques da Silva, J., 2018. Microbial extracellular polymeric substances improve water retention in dryland biological soil crusts. Soil Biology and Biochemistry 116, 67–69. URL: https://www.sciencedirect.com/science/article/pii/S003807171730593X, doi:10.1016/j.soilbio.2017.10.002.

Ahmed, M.A., Holz, M., Woche, S.K., Bachmann, J., Carminati, A., 2015. Effect of soil drying on mucilage exudation and its water repellency: a new method to collect mucilage. Journal of Plant Nutrition and Soil Science 178, 821–824. URL: https://onlinelibrary.wiley.com/doi/abs/10.1002/jpln.201500177, doi:10.1002/jpln.201500177. _eprint: https://onlinelibrary.wiley.com/doi/pdf/10.1002/jpln.201500177.

Ahmed, M.A., Kroener, E., Benard, P., Zarebanadkouki, M., Kaestner, A., Carminati, A., 2016. Drying of mucilage causes water repellency in the rhizosphere of maize: measurements and modelling. Plant and Soil 407, 161–171. URL: https://www.jstor.org/stable/44136917. publisher: Springer.

Ahmed, M.A., Sanaullah, M., Blagodatskaya, E., Mason-Jones, K., Jawad, H., Kuzyakov, Y., Dippold, M.A., 2018. Soil microorganisms exhibit enzymatic and priming response to root mucilage under drought. Soil Biology and Biochemistry 116, 410–418. URL: https://www.sciencedirect.com/science/article/pii/S0038071717304534, doi:10.1016/j.soilbio.2017.10.041.

Allison, S.D., Jastrow, J.D., 2006. Activities of extracellular enzymes in physically isolated fractions of restored grassland soils. Soil Biology and Biochemistry 38, 3245–3256. URL: https://www.sciencedirect.com/science/article/pii/S0038071706001878, doi:10.1016/j.soilbio.2006.04.011.

Benard, P., Zarebanadkouki, M., Hedwig, C., Holz, M., Ahmed, M., Carminati, A., 2018. Pore-Scale Distribution of Mucilage Affecting Water Repellency in the Rhizosphere. Vadose Zone Journal 17, 170013. URL: https://onlinelibrary.wiley.com/doi/abs/10.2136/vzj2017.01.0013, doi:10.2136/vzj2017.01.0013. _eprint: https://onlinelibrary.wiley.com/doi/pdf/10.2136/vzj2017.01.0013.

Bogati, K., Walczak, M., 2022. The Impact of Drought Stress on Soil Microbial Community, Enzyme Activities and Plants. Agronomy 12, 189. URL: https://www.mdpi.com/2073-4395/12/1/189, doi:10.3390/agronomy12010189. publisher: Multidisciplinary Digital Publishing Institute.

Bore, E.K., Halicki, S., Kuzyakov, Y., Dippold, M.A., 2019. Structural and physiological adaptations of soil microorganisms to freezing revealed by position-specific labeling and compound-specific 13C analysis. Biogeochemistry 143, 207–219. URL: 10.1007/s10533-019-00558-5, doi:10.1007/s10533-019-00558-5.

Burns, R.G., DeForest, J.L., Marxsen, J., Sinsabaugh, R.L., Stromberger, M.E., Wallenstein, M.D., Weintraub, M.N., Zoppini, A., 2013. Soil enzymes in a changing environment: Current knowledge and future directions. Soil Biology and Biochemistry 58, 216–234. URL: https://www.sciencedirect.com/science/article/pii/S0038071712004476, doi:10.1016/j.soilbio.2012.11.009.

Capitani, M.I., Ixtaina, V.Y., Nolasco, S.M., Tomás, M.C., 2013. Microstructure, chemical composition and mucilage exudation of chia (Salvia hispanica L.) nutlets from Argentina. Journal of the Science of Food and Agriculture 93, 3856–3862. doi:10.1002/jsfa.6327.

Carminati, A., Benard, P., Ahmed, M.A., Zarebanadkouki, M., 2017. Liquid bridges at the root-soil interface. Plant and Soil 417, 1–15. URL: 10.1007/s11104-017-3227-8, doi:10.1007/s11104-017-3227-8.

Chen, J., Qu, C., Lu, M., Zhang, M., Wu, Y., Gao, C., Huang, Q., Cai, P., 2023. Extracellular polymeric substances and mineral interfacial reactions control the simultaneous immobilization and reduction of arsenic (As(V)). Journal of Hazardous Materials 456, 131651. URL: https://www.sciencedirect.com/science/article/pii/S0304389423009342, doi:10.1016/j.jhazmat.2023.131651.

Cheng, C.H., Lehmann, J., Engelhard, M.H., 2008. Natural oxidation of black carbon in soils: Changes in molecular form and surface charge along a climosequence. Geochimica et Cosmochimica Acta 72, 1598– 1610. URL: https://www.sciencedirect.com/science/article/pii/S0016703708000306, doi:10.1016/j.gca.2008.01.010.

Cheng, H., Hill, P.W., Bastami, M.S., Jones, D.L., 2017. Biochar stimulates the decomposition of simple organic matter and suppresses the decomposition of complex organic matter in a sandy loam soil. GCB Bioenergy 9, 1110–1121. URL: https://onlinelibrary.wiley.com/doi/abs/10.1111/gcbb.12402, doi:10.1111/gcbb.12402. _eprint: https://onlinelibrary.wiley.com/doi/pdf/10.1111/gcbb.12402.

Chenu, C., 1993. Clay- or sand-polysaccharide associations as models for the interface between micro-organisms and soil: water related properties and microstructure. Geoderma 56, 143–156. URL: https://www.sciencedirect.com/science/article/pii/001670619390106U, doi:10.1016/0016-7061(93)90106-U.

Costa, O.Y.A., Raaijmakers, J.M., Kuramae, E.E., 2018. Microbial Extracellular Polymeric Substances: Ecological Function and Impact on Soil Aggregation. Frontiers in Microbiology 9. URL: https://www.frontiersin.org/journals/microbiology/articles/10.3389/fmicb.2018.01636/full, doi:10.3389/fmicb.2018.01636. publisher: Frontiers.

Crawley, M.J., 2013. The R book. 2nd ed ed., John Wiley & Sons Inc, Hoboken, N.J.

Dippold, M., Biryukov, M., Kuzyakov, Y., 2014. Sorption affects amino acid pathways in soil: Implications from position-specific labeling of alanine. Soil Biology and Biochemistry 72, 180–192. URL: https://www.sciencedirect.com/science/article/pii/S0038071714000169, doi:10.1016/j.soilbio.2014.01.015.

Fang, L., Huang, Q., Wei, X., Liang, W., Rong, X., Chen, W., Cai, P., 2010. Microcalorimetric and potentiometric titration studies on the adsorption of copper by extracellular polymeric substances (EPS), minerals and their composites. Bioresource Technology 101, 5774– 5779. URL: https://www.sciencedirect.com/science/article/pii/S096085241000386X, doi:10.1016/j.biortech.2010.02.075.

German, D.P., Weintraub, M.N., Grandy, A.S., Lauber, C.L., Rinkes, Z.L., Allison, S.D., 2011. Optimization of hydrolytic and oxidative enzyme methods for ecosystem studies. Soil Biology and Biochemistry 43, 1387– 1397. URL: https://www.sciencedirect.com/science/article/pii/S0038071711001301, doi:10.1016/j.soilbio.2011.03.017.

Gomez, J.D., Denef, K., Stewart, C.E., Zheng, J., Cotrufo, M.F., 2014. Biochar addition rate influences soil microbial abundance and activity in temperate soils. European Journal of Soil Science 65, 28–39. URL: https://onlinelibrary.wiley.com/doi/abs/10.1111/ejss.12097, doi:10.1111/ejss.12097. _eprint: https://onlinelibrary.wiley.com/doi/pdf/10.1111/ejss.12097.

Hagemann, N., Joseph, S., Schmidt, H.P., Kammann, C., Harter, J., Young, R., Varga, K., Tahery, S., Elliott, K., Mckenna, A., Albu, M., Mayrhofer, C., Obst, M., Conte, P., Dieguez Alonso, A., Orsetti, S., Subdiaga, E., Behrens, S., Kappler, A., 2017. Organic coating on biochar explains its nutrient retention and stimulation of soil fertility. Nature Communications 8, 1089. doi:10.1038/s41467-017-01123-0.

Holz, M., Leue, M., Ahmed, M.A., Benard, P., Gerke, H.H., Carminati, A., 2018. Spatial Distribution of Mucilage in the Rhizosphere Measured With Infrared Spectroscopy. Frontiers in Environmental Science 6. URL: https://www.frontiersin.org/journals/environmental-science/articles/10.3389/fenvs.2018.00087/full, doi:10.3389/fenvs.2018.00087. publisher: Frontiers.

Janu, R., Mrlik, V., Ribitsch, D., Hofman, J., Sedláček, P., Bielská, L., Soja, G., 2021. Biochar surface functional groups as affected by biomass feedstock, biochar composition and pyrolysis temperature. Carbon Resources Conversion 4, 36–46. URL: https://www.sciencedirect.com/science/article/pii/S2588913321000119, doi:10.1016/j.crcon.2021.01.003.

Joseph, S., Cowie, A.L., Van Zwieten, L., Bolan, N., Budai, A., Buss, W., Cayuela, M.L., Graber, E.R., Ippolito, J.A., Kuzyakov, Y., Luo, Y., Ok, Y.S., Palansooriya, K.N., Shepherd, J., Stephens, S., Weng, Z.H., Lehmann, J., 2021. How biochar works, and when it doesn’t: A review of mechanisms controlling soil and plant responses to biochar. GCB Bioenergy 13, 1731–1764. URL: https://onlinelibrary.wiley.com/doi/abs/10.1111/gcbb.12885, doi:10.1111/gcbb.12885. _eprint: https://onlinelibrary.wiley.com/doi/pdf/10.1111/gcbb.12885.

Kayoumu, M., Wang, H., Duan, G., 2025. Interactions between microbial extracellular polymeric substances and biochar, and their potential applications: a review. Biochar 7, 62. URL: 10.1007/s42773-025-00452-4, doi:10.1007/s42773-025-00452-4.

Kidinda, L.K., Babin, D., Doetterl, S., Kalbitz, K., Mujinya, B.B., Vogel, C., 2023. Extracellular polymeric substances are closely related to land cover, microbial communities, and enzyme activity in tropical soils. Soil Biology and Biochemistry 187, 109221. URL: https://www.sciencedirect.com/science/article/pii/S0038071723002833, doi:10.1016/j.soilbio.2023.109221.

Kohlmann, S., Greenberg, I., Joergensen, R.G., Dippold, M.A., Ludwig, B., 2025. Effects of low molecular weight organic substances, biochars and temperature on the microbial carbon use efficiency. Pedosphere URL: https://www.sciencedirect.com/science/article/pii/S1002016025000104, doi:10.1016/j.pedsph.2025.01.009.

Kuzyakov, Y., Subbotina, I., Chen, H., Bogomolova, I., Xu, X., 2009. Black carbon decomposition and incorporation into soil microbial biomass estimated by 14C labeling. Soil Biology and Biochemistry 41, 210–219. URL: https://www.sciencedirect.com/science/article/pii/S0038071708003544, doi:10.1016/j.soilbio.2008.10.016.

Lehmann, J., Kleber, M., 2015. The contentious nature of soil organic matter. Nature 528, 60–68. URL: https://www.nature.com/articles/nature16069, doi:10.1038/nature16069.

Lehmann, J., Rillig, M.C., Thies, J., Masiello, C.A., Hockaday, W.C., Crowley, D., 2011. Biochar effects on soil biota – A review. Soil Biology and Biochemistry 43, 1812–1836. URL: https://www.sciencedirect.com/science/article/pii/S0038071711001805, doi:10.1016/j.soilbio.2011.04.022.

Leng, L., Xiong, Q., Yang, L., Li, H., Zhou, Y., Zhang, W., Jiang, S., Li, H., Huang, H., 2021. An overview on engineering the surface area and porosity of biochar. Science of The Total Environment 763, 144204. URL: https://www.sciencedirect.com/science/article/pii/S0048969720377354, doi:10.1016/j.scitotenv.2020.144204.

McCully, M.E., Boyer, J.S., 1997. The expansion of maize rootcap mucilage during hydration. 3. Changes in water potential and water content. Physiologia Plantarum 99, 169–177. URL: https://onlinelibrary.wiley.com/doi/abs/10.1111/j.1399-3054.1997.tb03445.x, doi:10.1111/j.1399-3054.1997.tb03445.x. _eprint: https://onlinelibrary.wiley.com/doi/pdf/10.1111/j.1399-3054.1997.tb03445.x.

Mia, S., Dijkstra, F.A., Singh, B., 2017. Aging Induced Changes in Biochar’s Functionality and Adsorption Behavior for Phosphate and Ammonium. Environmental Science & Technology 51, 8359–8367. URL: 10.1021/acs.est.7b00647, doi:10.1021/acs.est.7b00647. publisher: American Chemical Society.

Munene, R., Mustafa, O., Loftus, S., Banfield, C.C., Rötter, R.P., Bore, E.K., Mweu, B., Mganga, K.Z., Otieno, D.O., Ahmed, M.A., Dippold, M.A., 2025. Contribution of arbuscular mycorrhiza and exoenzymes to nitrogen acquisition of sorghum under drought. Frontiers in Plant Science 16. URL: https://www.frontiersin.org/journals/plant-science/articles/10.3389/fpls.2025.1514416/full, doi:10.3389/fpls.2025.1514416. publisher: Frontiers.

Nazari, M., Bickel, S., Benard, P., Mason-Jones, K., Carminati, A., Dippold, M.A., 2022. Biogels in Soils: Plant Mucilage as a Biofilm Matrix That Shapes the Rhizosphere Microbial Habitat. Frontiers in Plant Science 12, 798992. URL: https://www.ncbi.nlm.nih.gov/pmc/articles/PMC8792611/, doi:10.3389/fpls.2021.798992.

Nazari, M., Bickel, S., Kuzyakov, Y., Bilyera, N., Zarebanadkouki, M., Wassermann, B., Dippold, M.A., 2024. Root mucilage nitrogen for rhizosphere microorganisms under drought. Biology and Fertility of Soils 60, 639–647. URL: https://link.springer.com/10.1007/s00374-024-01827-8, doi:10.1007/s00374-024-01827-8.

Niedeggen, D., Rüger, L., Oburger, E., Santangeli, M., Ahmed, M., Vetterlein, D., Blagodatsky, S., Bonkowski, M., 2024. Microbial utilisation of maize rhizodeposits applied to agricultural soil at a range of concentrations. European Journal of Soil Science 75, e13530. URL: https://onlinelibrary.wiley.com/doi/abs/10.1111/ejss.13530, doi:10.1111/ejss.13530. _eprint: https://bsssjournals.onlinelibrary.wiley.com/doi/pdf/10.1111/ejss.13530.

Okolo, C.C., Bore, E., Gebresamuel, G., Zenebe, A., Haile, M., Nwite, J.N., Dippold, M.A., 2022. Priming effect in semi-arid soils of northern Ethiopia under different land use types. Biogeochemistry 158, 383–403. URL: 10.1007/s10533-022-00905-z, doi:10.1007/s10533-022-00905-z.

Palansooriya, K.N., Wong, J.T.F., Hashimoto, Y., Huang, L., Rinklebe, J., Chang, S.X., Bolan, N., Wang, H., Ok, Y.S., 2019. Response of microbial communities to biochar-amended soils: a critical review. Biochar 1, 3–22. URL: 10.1007/s42773-019-00009-2, doi:10.1007/s42773-019-00009-2.

Piepho, H.P., 2009. Data Transformation in Statistical Analysis of Field Trials with Changing Treatment Variance. Agronomy Journal 101, 865–869. URL: https://onlinelibrary.wiley.com/doi/abs/10.2134/agronj2008.0226x, doi:10.2134/agronj2008.0226x. _eprint: https://acsess.onlinelibrary.wiley.com/doi/pdf/10.2134/agronj2008.0226x.

Razavi, B.S., Blagodatskaya, E., Kuzyakov, Y., 2015. Nonlinear temperature sensitivity of enzyme kinetics explains canceling effect—a case study on loamy haplic Luvisol. Frontiers in Microbiology 6. URL: https://www.frontiersin.orghttps://www.frontiersin.org/journals/microbiology/articles/10.3389/fmicb.2015.01126/full, doi:10.3389/fmicb.2015.01126. publisher: Frontiers.

Redmile-Gordon, M., Gregory, A.S., White, R.P., Watts, C.W., 2020. Soil organic carbon, extracellular polymeric substances (EPS), and soil structural stability as affected by previous and current land-use. Geoderma 363, 114143. URL: https://www.sciencedirect.com/science/article/pii/S0016706119312091, doi:10.1016/j.geoderma.2019.114143.

Sun, D., Meng, J., Xu, E.G., Chen, W., 2016. Microbial community structure and predicted bacterial metabolic functions in biochar pellets aged in soil after 34 months. Applied Soil Ecology 100, 135–143. URL: https://www.sciencedirect.com/science/article/pii/S0929139315301566, doi:10.1016/j.apsoil.2015.12.012.

Tang, J., Riley, W.J., 2019. A Theory of Effective Microbial Substrate Affinity Parameters in Variably Saturated Soils and an Example Application to Aerobic Soil Heterotrophic Respiration. Journal of Geophysical Research: Biogeosciences 124, 918–940. URL: https://onlinelibrary.wiley.com/doi/abs/10.1029/2018JG004779, doi:10.1029/2018JG004779. _eprint: https://agupubs.onlinelibrary.wiley.com/doi/pdf/10.1029/2018JG004779.

Team, R.C., 2024. R: A language and environment for statistical computing. URL: https://www.R-project.org/.

Vance, E.D., Brookes, P.C., Jenkinson, D.S., 1987. An extraction method for measuring soil microbial biomass C. Soil Biology and Biochemistry 19, 703–707. URL: https://www.sciencedirect.com/science/article/pii/0038071787900526, doi:10.1016/0038-0717(87)90052-6.

Wang, J., Xiong, Z., Kuzyakov, Y., 2016. Biochar stability in soil: meta-analysis of decomposition and priming effects. GCB Bioenergy 8, 512–523. URL: https://onlinelibrary.wiley.com/doi/abs/10.1111/gcbb.12266, doi:10.1111/gcbb.12266. _eprint: https://onlinelibrary.wiley.com/doi/pdf/10.1111/gcbb.12266.

Wu, J., Joergensen, R.G., Pommerening, B., Chaussod, R., Brookes, P.C., 1990. Measurement of soil microbial biomass C by fumigation-extraction—an automated procedure. Soil Biology and Biochemistry 22, 1167–1169. URL: https://www.sciencedirect.com/science/article/pii/0038071790900463, doi:10.1016/0038-0717(90)90046-3.

Zhang, P., Duan, W., Peng, H., Pan, B., Xing, B., 2021. Functional Biochar and Its Balanced Design. ACS Environmental Au 2, 115–127. URL: https://www.ncbi.nlm.nih.gov/pmc/articles/PMC10114722/, doi:10.1021/acsenvironau.1c00032.

Zimmerman, A.R., Gao, B., Ahn, M.Y., 2011. Positive and negative carbon mineralization priming effects among a variety of biochar-amended soils. Soil Biology and Biochemistry 43, 1169– 1179. URL: https://www.sciencedirect.com/science/article/pii/S0038071711000769, doi:10.1016/j.soilbio.2011.02.005.

