## Appendix1 for "Effect of biogel C addition on biochar degradation and microbial activities"

**Appendix 1: Summary of final ANOVA models after stepwise removal of non-significant terms and marginal means.**

| **Source** | **Degrees of freedom** | **Sum of squares** | **Mean square** | **F** | **P** |
| --- | --- | --- | --- | --- | --- |
| **Box-Cox-transformed Cumulative^14^CO_2_ efflux** |  |  |  |  |  |
| **Phase I** |  |  |  |  |  |
| Block | 3 | 1.0 | 0.3 | 0.9 | 0.47 |
| Biochar | 1 | 2.4 | 2.5 | 6.2 | 0.02 |
| Residuals | 27 | 10.3 | 0.4 |  |  |
| **Phase II** |  |  |  |  |  |
| Block | 3 | 0.4 | 0.1 | 0.4 | 0.79 |
| Biochar | 1 | 3.0 | 3.0 | 9.4 | 4.8*10^-3^ |
| Residuals | 27 | 8.5 | 0.3 |  |  |
| **Cumulative^14^CO_2_ efflux (sustained phase)**  **(% of ^14^C input)** |  |  |  |  |  |
| Block | 3 | 9.6 | 3.2 | 0.2 | 0.87 |
| Biochar | 1 | 80.5 | 80.5 | 6.1 | 0.02 |
| Residuals | 27 | 354.0 | 13.1 |  |  |
| **MBC (µg C g^-1^ soil)** |  |  |  |  |  |
| Block | 3 | 1.8*10^4^ | 5.9*10^3^ | 8.5 | 1.5*10^-4^ |
| Moisture | 1 | 5.5*10^3^ | 5.5*10^3^ | 7.9 | 7.6*10^-3^ |
| Residuals | 43 | 3.0*10^4^ | 7.0*10^2^ |  |  |
| **^14^C incorporation (MBC)**  **(µg C g^-1^ soil)** |  |  |  |  |  |
| Block | 3 | 0.5 | 0.2 | 0.5 | 0.70 |
| Moisture | 1 | 0.9 | 0.9 | 0.5 | 0.49 |
| Mucilage | 1 | 2.0 | 2.0 | 5.4 | 0.03 |
| Biochar | 1 | 0.5 | 0.5 | 1.2 | 0.28 |
| Moisture:Mucilage | 1 | 0.1 | 0.1 | 0.2 | 0.65 |
| Moisture:Biochar | 1 | 0.1 | 0.1 | 0.4 | 0.59 |
| Mucilage:Biochar | 1 | 0.4 | 0.4 | 1.0 | 0.32 |
| Moisture:Mucilage:Biochar | 1 | 3.3 | 3.3 | 9.0 | 6.9*10^-3^ |
| Residuals | 21 | 7.8 | 0.4 |  |  |
| **β-glucosidase** |  |  |  |  |  |
| **Box-Cox-transformed V_max_** |  |  |  |  |  |
| Block | 3 | 2.9*10^8^ | 9.6*10^7^ | 1.5 | 0.24 |
| Moisture | 1 | 3.6*10^8^ | 3.6*10^8^ | 5.1 | 0.02 |
| Residuals | 43 | 2.8*10^9^ | 6.6*10^7^ |  |  |
| **K_m_ (µM g^-1^)** |  |  |  |  |  |
| Block | 3 | 2.3*10^4^ | 7.8*10^3^ | 6.7 | 1.1*10^-3^ |
| Moisture | 1 | 6.5*10^3^ | 6.5*10^3^ | 5.6 | 0.02 |
| Mucilage | 1 | 6.8*10^3^ | 6.8*10^3^ | 5.8 | 0.02 |
| Biochar | 2 | 5.2*10^3^ | 2.6*10^3^ | 2.2 | 0.12 |
| Moisture:Mucilage | 1 | 2.7*10^4^ | 2.7*10^4^ | 23.4 | 2.6*10^-5^ |
| Moisture:Biochar | 2 | 1.1*10^4^ | 5.7*10^3^ | 5.0 | 0.01 |
| Mucilage:Biochar | 2 | 8.0*10^3^ | 4.0*10^3^ | 3.5 | 0.04 |
| Residuals | 35 | 4.0*10^4^ | 1.2*10^3^ |  |  |
| **Box-Cox-transformed Peroxidase activity** |  |  |  |  |  |
| Block | 3 | 0.0 | 0.0 | 0.1 | 0.93 |
| Moisture | 1 | 0.0 | 0.0 | 4.0 | 0.06 |
| Mucilage | 1 | 0.0 | 0.0 | 1.4 | 0.25 |
| Biochar | 2 | 0.2 | 0.1 | 13.0 | 6.9*10^-5^ |
| Moisture:Mucilage | 1 | 0.0 | 0.0 | 3.7 | 0.06 |
| Moisture:Biochar | 2 | 0.0 | 0.0 | 2.0 | 0.16 |
| Mucilage:Biochar | 2 | 1.0 | 0.0 | 4.0 | 0.03 |
| Moisture:Mucilage:Biochar | 2 | 1.0 | 0.0 | 5.0 | 0.01 |
| Residuals | 33 | 0.3 | 0.0 |  |  |
| **Box-Cox-transformed BET** |  |  |  |  |  |
| Block | 3 | 189.5 | 63.2 | 279.6 | 6.8*10^-16^ |
| Days | 1 | 22.6 | 22.6 | 99.9 | 5.3*10^-9^ |
| Residuals | 19 | 4.3 | 0.2 |  |  |
| **Marginal means** |  |  |  |  |  |
| **Cumulative ^14^CO_2_ efflux**  **(% of ^14^C input)** |  | | | | |
| **Phase I** | Means^b^ for aged and fresh biochar 2.9 (2.3, 3.6), 4.2 (3.3, 5.1) | | | | |
| **Phase II** | Means^b^ for aged and fresh biochar 14.4 (12.6, 16.4), 18.9 (16.6, 21.3) | | | | |
| **Cumulative ^14^CO_2_ efflux (Sustained phase)**  **(% of ^14^C input)** | Means^a^ for aged and fresh biochar 11.6, 14.8 (1.28) | | | | |
| **MBC (µg C g^-1^ soil)** | Means^a^ for 30% and 70% WHC 129.0, 150.4 (5.4) | | | | |
| **ꞵ-glucosidase** |  | | | | |
| **V_max_ (nM g^-1^ h^-1^)** | Means^b^ for 30% and 70% WHC 600.9 (497.0, 696.6), 751.9 (661.6, 836.7) | | | | |
| **BET (m^2^/g)** | Means^b^ for day 1 and day 85 10.0 (9.7, 10.3), 11.9 (11.7, 12.2) | | | | |
| ^a^Values in the parentheses gives the standard errors for the differences of means.  ^b^Predicted marginal means with confidence limits in the parentheses were back-transformed and may be interpreted as medians and confidence limits of the medians (Piepho, 2009).  Marginal means were not calculated for ^14^C Incorporation (MBC), K_m_ and peroxidase activity. | | | | | |
